# Constitutive expression of full-length or partial of *SOC1* genes for yield enhancement in tomato

**DOI:** 10.1101/2025.06.03.657751

**Authors:** Gharbia H. Danial, Jirapa Jaikham, Guo-qing Song

## Abstract

Manipulating the expression of flowering pathway genes holds potential for regulating tomato fruit productivity. *SUPPRESSOR OF OVEREXPRESSION OF CONSTANS 1* (*SOC1*) is a MADS-box gene that serves as a key integrator in the flowering pathway. In this study, two full-length *SOC1* genes cloned from maize (*ZmSOC1*) and soybean (*GmSOC1*), along with a partial *SOC1* gene from blueberry (*VcSOC1K*, containing the K-domain), were individually transformed into tomato for constitutive expression. Phenotypically, the expression of *VcSOC1K* and *ZmSOC1*, but not *GmSOC1*, led to early flowering. Most transgenic lines for all three constructs exhibited a significant increase in fruit number per plant. More importantly, compared to non-transgenic plants, all three constructs resulted in varying degrees of increased fruit production per plant, primarily through enhanced branching. At the transcriptomic level, comparative analysis of *GmSOC1* revealed the broader impact of the transformed genes. The increased expression of *CLF* and *EZA1* appears to explain the unchanged flowering time of the *GmSOC1* transgenic plants, while the repressed expression of *DWARF* genes likely contributes to enhanced branching. Additionally, numerous genes associated with biotic and abiotic stress tolerance displayed differential expression. These findings demonstrate that constitutive expression of either full-length or partial *SOC1* has the potential to enhance tomato fruit production by modulating multiple pathways, at least at the transcript levels.

## Introduction

The MADS-box gene family encodes transcription factors that are present in all eukaryotic organisms, playing essential roles in animals, plants, and fungi ^1–3^. In plants, these genes are crucial for a wide range of physiological and developmental processes ^4^. While MADS-box proteins have been extensively studied for their roles in regulating flower development, emerging research indicates their involvement in fruit development, embryo establishment, vegetative organ development, and stress resistance. These findings highlight the significance and functional diversity of this gene superfamily in plant development ^5–7^.

The plant MADS-box family includes type I and II genes ^8,9^. Type I MADS box factors are crucial regulators of plant reproduction; they play key roles in the development of the female gametophyte, embryo, and endosperm development ^8^. Type II genes have been extensively studied and further classified into MIKC^C^- and MIKC*-type two subgroups ^10^. MIKC^C^-type genes are the most thoroughly investigated members of the plant MADS-box family and encode proteins containing four distinct domains: M, I, K, and C. The M (DNA-binding) domain is the most conserved and is shared across all MADS-box genes. The I (intervening) domain, while less conserved, facilitates the specificity of DNA-binding dimer formation. The K (keratin-like) domain is structurally conserved and mediates protein–protein interactions, while the C (C-terminal) domain, the least conserved, is involved in ternary complex formation and transcriptional activation ^10^. MIKC*-type genes are believed to have originated from ancestral MIKC^C^-type genes through either elongation of the I region or truncation of the K box ^10–12^. These genes are crucial for pollen development ^12,13^. This structure-function relationship highlights the critical role of MADS-box genes in regulating diverse developmental processes, including fruit development, embryogenesis, vegetative organ formation, and stress responses.

MIKC^C^-type MADS-box (MIKC^C^) contribute floral organogenesis and flowering transition^14–16^. Among these, *SUPPRESSOR OF OVEREXPRESSION OF CONSTANS 1* (*SOC1*) serves as a conserved floral activator and integrator within the plant flowering pathway ^17–20^. Given the multifunctional roles of the SOC1 protein in regulating plant development, its expression can be modulated through genetic engineering to potentially enhance crop yield ^21–25^.

Crop yield, such as in tomato, is influenced by genetic background, environmental conditions, and management practices. The potential for crop yield is genetically determined through the interactions of multiple genes and complex gene networks ^26^. Strategically, genetic improvement can be achieved by incorporating genes for biotic resistance and abiotic stress tolerance through breeding to protect yields, while modifying key genes related to growth and development can further enhance productivity ^27^.

Tomato (*Solanum lycopersicum* L.) is an important vegetable crop and a valuable source of essential nutrients and phytochemicals, including vitamin C, lycopene, and antioxidants ^28^. Enhancing yield is a top breeding priority for fresh-market tomato ^26^. This can be achieved through either traditional breeding or genetic engineering ^29,30^. However, since yield is a trait controlled by multiple genes, both approaches present significant challenges. Although numerous genes have been functionally characterized in tomato, only a few, particularly those involved in flowering pathways, including four tomato *SOC1* genes, have shown potential for increasing yield ^29–32^. Yet the specific effects of these genes on yield remain unexplored.

Due to concerns that overexpression of the tomato *SOC1* gene may lead to dosage-related effects, this study investigated the constitutive expression of two heterologous *SOC1* genes, *GmSOC1* from soybean (*Glycine max*) and *ZmSOC1* from maize (*Zea mays*), along with a truncated derivative of blueberry (*Vaccinium corymbosum*) *SOC1* lacking the M domain (*VcSOC1K*), to assess their potential for improving tomato yield.

## Results

### Phylogenetic analysis of tomato’s SOC1 and SOC1-like proteins

The tomato proteome in Phytozome v13 includes one SOC1 protein and four SOC1-like proteins, which have been identified as SlTM3, SlSTM3, SlMBP18, and SlMBP23 by Zahn et al ^32^. These five genes are distributed across four chromosomes (Fig. 1A). Their protein sequences exhibit high similarity to *Arabidopsis* SOC1, GmSOC1, ZmSOC1, and VcSOC1K. Notably, GmSOC1 and ZmSOC1 cluster with SlMBP18 and the SlSOC1-like protein XP_004252597.2, while VcSOC1 clusters with SlTM3, SlSTM3, SlMBP23, and *Arabidopsis* SOC1.

**Fig. 1.**
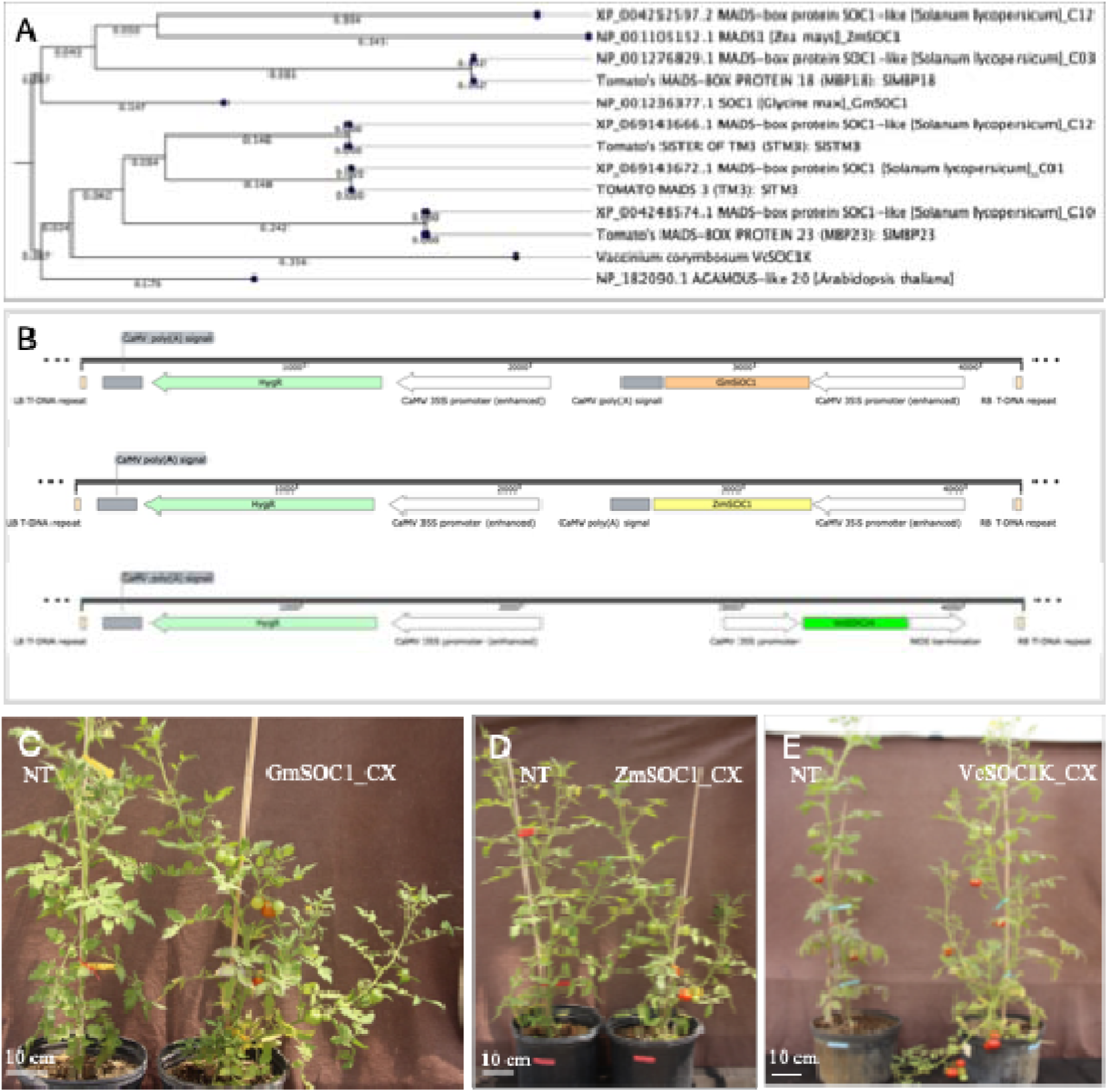
Genes and constructs used for generating transgenic tomato. **(A)** Phylogenetic analysis of five tomato SOC1 and SOC1-like proteins from four chromosomes. Protein sequences of TOMATO MADS3 (SlTM3), SISTER OF TM3 (SlSTM3), SlMBP18. and SlMBP23 are from Zahn et al., 2023 ^32^. C01, C03, C10, and C12 are chromosome numbers. The analysis reveals similarities to *Arabidopsis* SOC1, soybean GmSOC1, maize ZmSOC1, and blueberry SOC1 K-domain (VcSOC1K), using the Neighbor Joining method with the Jukes-Cantor protein distance metric and 500 bootstrap replicates. **(B)** T-DNA regions of three pCAMBIA1300-derived constructs containing *GmSOC1*, *ZmSOC1*, and *VcSOC1K* genes. HygR: the hygromycin B resistance gene. LB and RB: the left and right borders, respectively. **(C)** Constitutive expression of *GmSOC1* (GmSOC1-CX) in T_0_ plants and non-transgenic (NT) regenerants. **(D)** Constitutive expression of *ZmSOC1* (ZmSOC1-CX) in T_0_ plants and NT regenerants. **(E)** Constitutive expression of *VcSOC1K* (VcSOC1K-CX) in T_0_ plants and NT regenerants.

In the transcriptome reference assembled from all transcripts in the leaves of the ’Ailsa Craig’ cultivar used in this study, transcripts corresponding to SlTM3, SlSTM3, SlMBP18, and the protein identical to XP_004252597.2 were detected, while SlMBP23 was absent.

The conserved nature of *SOC1* and *SOC1*-like genes suggests that their orthologs *SlTM3*, *SlSTM3*, *SlMBP18*, and *SlMBP23* in tomato may play similar roles to those in other plant species. Consequently, the constitutive expression of *GmSOC1*, *ZmSOC1*, and *VcSOC1K* may have the potential to influence fruit production.

### Phenotypic changes in T_0_ transgenic plants

A total of 69 hygromycin-resistant T_0_ transgenic lines for GmSOC1_CX, 51 for ZmSOC1_CX, and 92 for VcSOC1K_CX were obtained across the three constructs, providing sufficient material for selecting comparable lines for phenotyping and seed harvesting (Fig. 1B-E).

The constitutive expression of *GmSOC1* (GmSOC1_CX) resulted in non-significant changes, such as earlier flowering and increases in plant height and the number of flower clusters (Supplementary Fig. S1A-C). Significant changes included increases in branch and fruit numbers, enhanced fruit production, larger fruit size, and earlier fruit maturation (Supplementary Fig. S1D-H).

The constitutive expression of *ZmSOC1* (ZmSOC1_CX) led to significant changes, including earlier flowering, earlier fruit maturation, and increased fruit number and production (Supplementary Fig. S1A, E-G). Non-significant changes included decreases in plant height, numbers of flower clusters and branches, and fruit size (Supplementary Fig. S1B-D, H).

The constitutive expression of *VcSOC1K* (VcSOC1K_CX) induced significant changes, including earlier flowering, increased numbers of flower clusters and fruits, enhanced fruit production, and reduced fruit size (Supplementary Fig. S1A, C, F-H). It also caused non-significant changes, such as earlier fruit maturation and increases in plant height and branch number (Supplementary Fig. S1B, D, E).

Taken together, GmSOC1_CX, ZmSOC1_CX, and VcSOC1K_CX all contributed to increased fruit numbers and production (Supplementary Fig. S1).

### Phenotypic changes in T_1_ transgenic plants

T_1_ transgenic plants were phenotypically compared with their corresponding plants (Fig. 2A-C). Under greenhouse conditions, among the nine groups with constitutive expression, ZmSOC1_CX17 was the only group of T_1_ transgenic plants to exhibit significantly earlier flowering, while the other T_1_ groups showed no difference in flowering time (Fig. 2D).

**Fig. 2.**
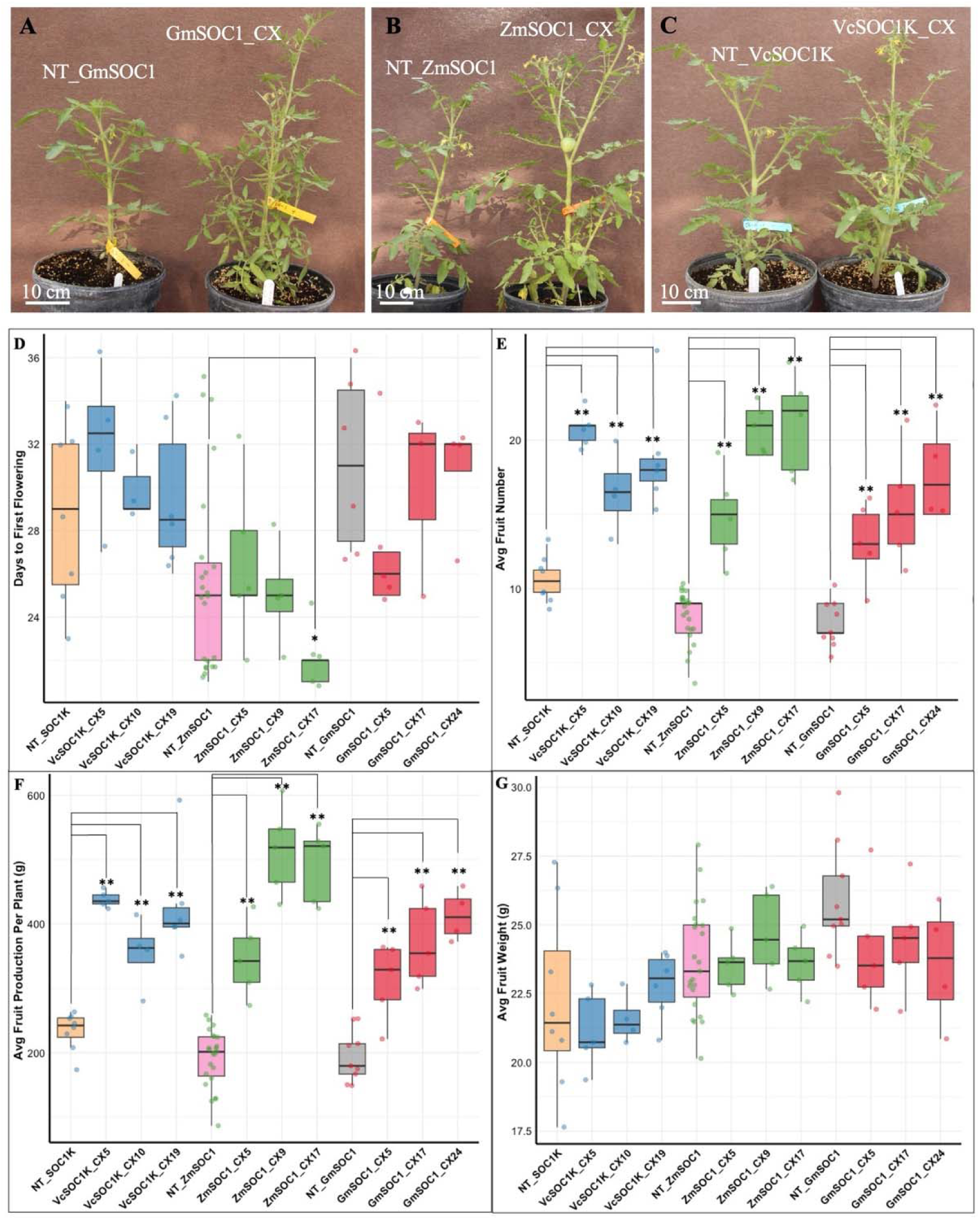
Phenotypic comparisons of three T_1_ transgenic lines for GmSOC1_CX, ZmSOC1_CX, and VcSOC1K_CX, alongside their corresponding non-transgenic (NT) seedlings (NT_GmSOC1, NT_ZmSOC1, and NT_VcSOC1K), with *n* > 3 for each transgenic and NT group. **(A-C)** Representative T_1_ plants. **(D)** Days to the appearance of the first flower after transplantation into a one-gallon pot. **(E)** Total number of fruits harvested. **(F)** Total weight of harvested fruits. **(G)** Average weight per fruit. The y-axis shows averages, and bars indicate standard deviations. Significance codes: \*\**p* < 0.01, \**p* < 0.05.

Consistent with observations in T_0_ plants, GmSOC1_CX, ZmSOC1_CX, and VcSOC1K_CX all resulted in increased fruit numbers and overall production, while the average fruit size showed no significant decrease (Fig. 2E-G).

### Transcriptomic analysis of T_1_ GmSOC1_CX plants

The T_1_ GmSOC1_CX plants were selected for transcriptome analysis to confirm the presence of transgenes and identify genes responsive to *GmSOC1* expression in tomato. This selection builds on previous transcriptome analyses of ZmSOC1_CX in soybean and maize, as well as VcSOC1K_CX in blueberry and maize, which have been reported ^22–24,33^.

The assembled transcriptome reference comprised 61,563 transcripts corresponding to 20,455 annotated genes. In GmSOC1_CX leaves, 565 differentially expressed transcripts (DETs) associated with 479 differentially expressed genes (DEGs) were identified. Among these DEGs, the two transgenes, *GmSOC1* and hygromycin phosphotransferase (*hph*), showed fold change (FC) of 23,170 [Log_2_^FC(transgenic/non-transgenic)^ = 14.5] and 7,131 [Log_2_^FC(transgenic/non-transgenic)^ = 12.8], respectively, confirming the expression of *GmSOC1* (Table 1). Additionally, RT-qPCR analysis of six selected DETs yielded results consistent with their RNA-seq data (Fig. 3), supporting the reliability of the RNA-seq findings.

**Fig. 3.**
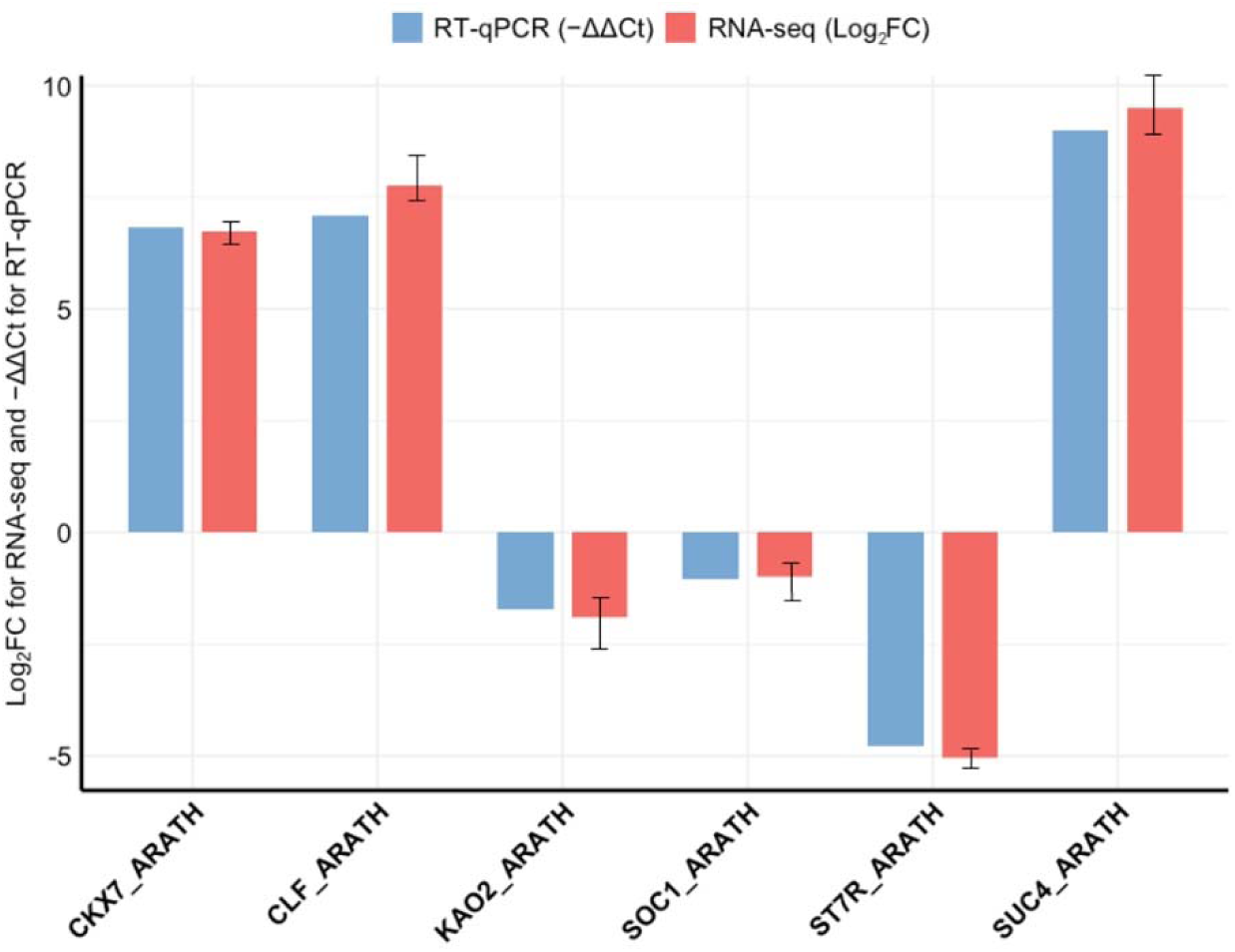
RT-qPCR analysis of the six selected DEGs listed in the Table 1. The −ΔΔCt values represent the average of three biological replicates and three technical replicates for each DEG. Tomato *ACTIN* gene SlACTIN12 (TRINITY_DN938_c0_g2_i1: ACT12_SOL) was used as the reference gene for normalization. Error bars represent standard deviation.

**Table 1.**
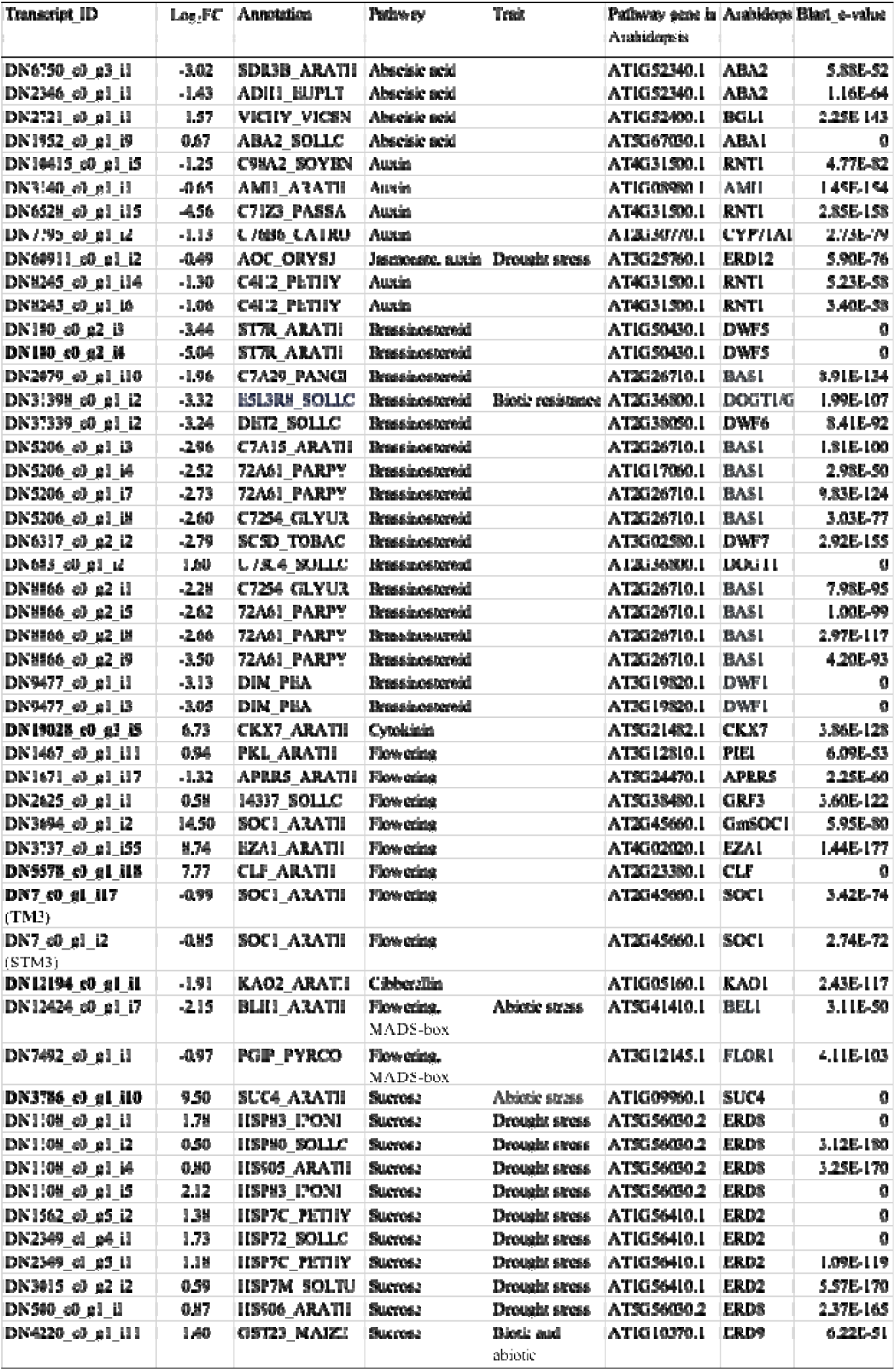
Differentially expressed transcripts (DETs) of flowering pathway, hormone, MADS-box genes, and sucrose-related genes in leaves of GmSOC1-CX plants. Log_2_FC: Log_2_(Fold change) = Log_2_(transgenic/non-transgenic). The transcript DN36964_c0_g1_i2 corresponds to the transformed *GmSOC1* gene from soybean. The bold DETs were further verified by RT-qPCR.

Analysis of the 479 DEGs using the GOSlim_Plants ontology file in BiNGO identified 27 overrepresented Gene Ontology (GO) terms. These included 11 terms under "Biological Process", seven under "Molecular Function", and nine under "Cellular Component" (Fig. 4). These overrepresented GO terms suggest a broad impact of GmSOC1_CX on plant development at the transcript level, contributing to the phenotypic changes observed in GmSOC1_CX plants.

**Fig. 4.**
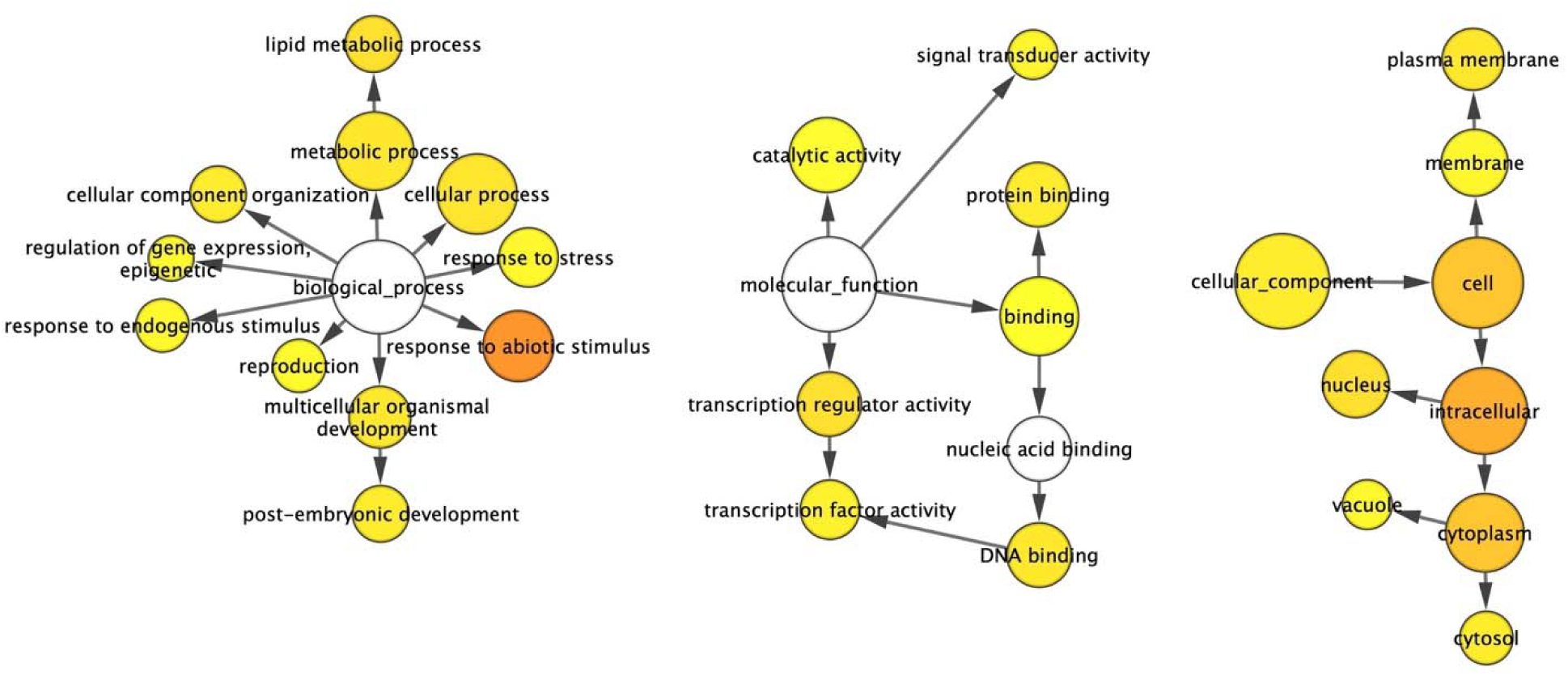
Gene networks of differentially expressed transcripts (DETs) identified from the comparison between the GmSOC1_CX and non-transgenic leaves. The ontology file of GOSlim_Plants in BiNGO was used to identify overrepresented GO terms (*p* < 0.05). Bubble size and color indicate the frequency of the GO term and the P-value, respectively.

For example, six overrepresented GO terms within the “Biological Process” category, namely “reproduction”, “response to abiotic stimulus”, “response to stress”, “post-embryogenic development”, “regulation of gene expression, epigenetic”, and “response to endogenous stimulus”, affect fruit production. These processes are linked through the overrepresented GO terms in the “Molecular Function” category, which regulate “transcription regulator activity” and “catalytic activity” via “binding” (Fig. 4).

Further analysis of the 565 DETs for key genes identified 51 key DETs associated with flowering, phytohormones, and sucrose. These were annotated to 39 DEGs and exhibited high similarity to 28 *Arabidopsis* genes (Table 1). These DETs were annotated to 39 DEGs and showed high similarities to 28 *Arabidopsis* genes (Table 1). Among the 39 DEGs, *GmSOC1* was highly expressed (Log_2_^FC(transgenic/non-transgenic)^ = 14.5) and suppressed the expression of two major endogenous tomato *SOC1* orthologues, *SlTM3* (TRINITY_DN7_c0_g1_i17) and *SlSTM3* (TRINITY_DN7_c0_g1_i2). In tomato, *SlTM3* and *SlSTM3* promote the floral transition but serve opposing roles in inflorescence development ^32^. High expression of *SLSTM* genes contributes to highly branched inflorescences ^31^.

The transcription factor BEL1-like homeodomain protein 1 (*BLH1*), which shares high similarity with the MADS-box gene *BEL1*, was downregulated. BLH family transcription factors are multifunctional, playing critical roles in plant morphogenesis, flower and fruit development, and responses to various environmental factors ^34^. Similarly, the polygalacturonase inhibitor precursor (*PGIP*), which exhibits a high similarity to the MADS-box gene *FLOR1*, known to promote flowering under long-day conditions (Torti et al., 2012), was also repressed.

The expression of two Histone-lysine N-methyltransferase genes *CURLY LEAF* (*CLF*) and *ENHANCER OF ZESTE 1 POLYCOMB REPRESSIVE COMPLEX 2 SUBUNIT* (*EZA1*), was enhanced. These genes encode catalytic subunits of the polycomb group (PcG) multiprotein complex. *CLF* is essential for regulating floral development by repressing the AGAMOUS homeotic gene. It achieves this by forming a nuclear complex with *EZA1* and other components, targeting ABA- and glucose-responsive elements ^35–38^. Notably, a loss-of-function mutant of *Brassica rapa* exhibits early flowering ^39^, suggesting that the upregulation of *CLF* and *EZA1* may contribute to either no significant impact or delaying flowering (Fig. 5).

**Fig. 5.**
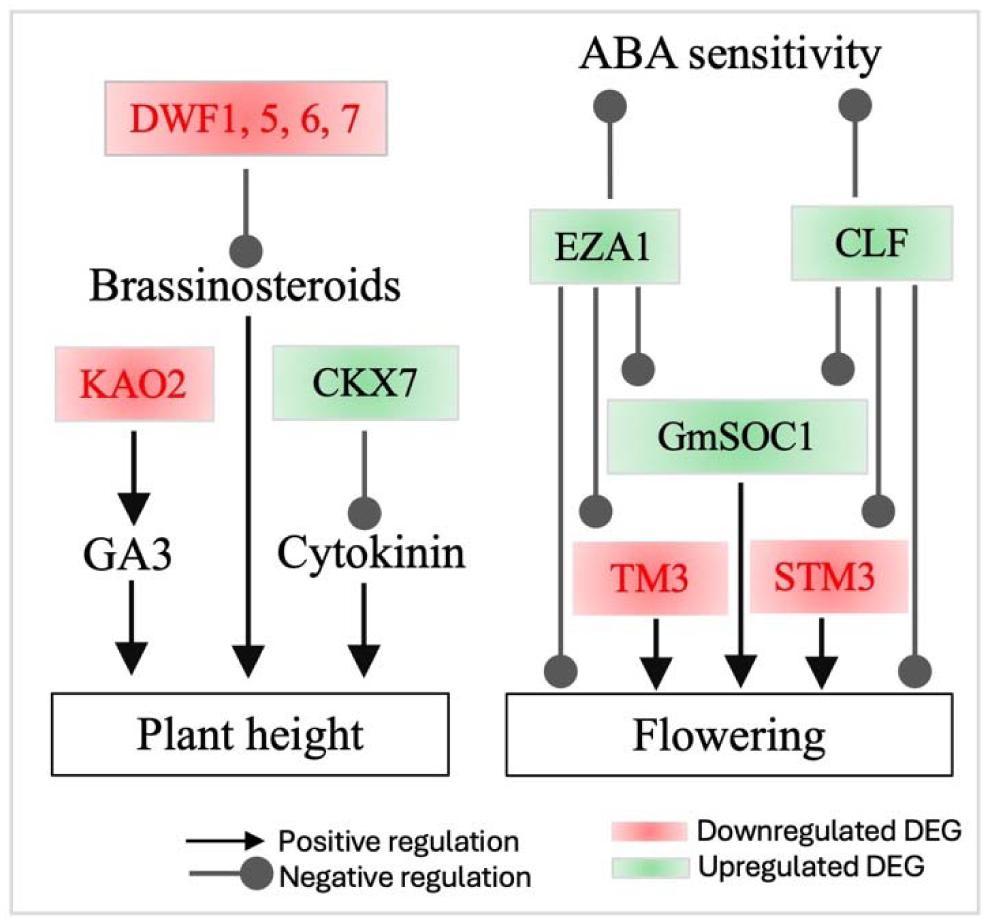
Interactions among major differentially expressed genes (DEGs) influencing tomato flowering and plant height in the leaves of GmSOC1_CX. The DEG data are presented in Table 1, and the proposed interactions are based on published information.

Of the genes involved in brassinosteroids (BRs) metabolism ^40^, the DEGs with repressed expressions included *BAS1* and four *DWARF* genes (*DWF1*, *DWF5*, *DWF6*, and *DWF7*) (Table 1); these DEG can functionally lead to BRs-deficient changes, which are often associated with decreased BRs that can affect multiple agronomic traits ^41–43^. The E5L3R8_SOLLC encodes tomato GLYCOALKALOID METABOLISM1 (GAME1) involved in the steroidal alkaloids (SAs) pathway ^44,45^. The E5L3R8_SOLLC showed decreased expression (Table 1), suggesting a potential lower production of α-tomatine that has impact on plant defense system and fruit quality ^44,46,47^.

For the cytokinin pathway genes, the expression of *CKX7* was enhanced, meaning a potential of a reduced cytokinin level in the GmSOC1-CX plants ^48^. Similarly, a decreased expression of the *KAO2* was found (Table 1), indicating a likely lower level of GA production ^49,50^.

Among the DEGs related to sucrose, the tomato sucrose transporter *SlSUT4* (*SUC4_ARATH*) was upregulated (Table 1). While the downregulation of *SlSUT4* has been shown to promote flowering by enhancing sucrose transport to the shoot apex, its overexpression does not significantly impact flowering time or the expression of key genes in the flowering pathway ^51^.

Additionally, seven DEGs of HEAT SHOCK PROTEINs showed upregulation (Table 1), potentially contributing to enhanced tolerance to various abiotic stresses ^52^. Similarly, the increased expression of the *GLUTATHIONE S-TRANSFERASE (GST) 23* is associated with both biotic and abiotic resistance ^53^. In contrast, the tomato ALCOHOL DEHYDROGENASE

(ADH1_EUPLT) and its homolog YFE37, annotated as SDR3B_ARATH, were both repressed. This reduction in expression may negatively impact disease resistance ^54^.

## Discussion

Tomato is commonly used as a model plant to study fruit-related traits, primarily due to its suitability for genetic transformation. In this study, we demonstrate for the first time that the ectopic expression of full-length or partial *SOC1* can enhance tomato fruit production per plant through a complex interaction of multiple genes and pathways. It is worth noting that ’Ailsa Craig’ is an indeterminate tomato variety, which adds complexity to phenotyping flowering and fruiting traits in this study. When T_0_ plants were examined, they were of similar size at the time of re-potting. However, for T_1_ plants, which were grown from seed germination, plant sizes were more variable at the time of re-potting. Importantly, phenotyping was conducted prior to genotyping by PCR, which minimized potential bias in the phenotypic data and ensured its reliability.

### Flowering time and the expression of Full-length *SOC1* and partial *SOC1*

As a key integrator in the plant flowering pathway ^17,55,56^, enhanced expression of *SOC1* or its orthologues, either though overexpression or ectopic expression, can promote flowering. This phenomenon has been reported for many plant species, including tomato ^32^. Among the five *SOC1* and *SOC1-like* genes identified in tomato (Fig. 1), *SlTM3* and *SlSTM3* play a more significant role in flowering initiation compared to *SlMBP23* and *SlMBP18*, at least in the indeterminate cultivar Moneyberg ^32^. Additionally, the high expression of *SlSTM3* has been linked to a highly branched inflorescence phenotype ^31^.

In this study, phylogenetic analysis revealed that GmSOC1 and ZmSOC1 clustered closely with SlMBP18 and the fifth SlSOC1-like gene, while VcSOC1K showed closer similarity to SlTM3, SlSTM3, and SlMBP23 (Fig. 1). Phenotypic observations demonstrated that both ZmSOC1-CX and VcSOC1-CX transgenic plants exhibited earlier flowering compared to non-transgenic controls, with statistically significant differences (*P* < 0.01) in the T_0_ generation and no significant differences (*P* < 0.01) in the T_1_ generation (Fig. 2D and Supplementary Fig. S1A). In contrast, GmSOC1-CX plants did not show significant changes in flowering time across both T0 and T_1_ generations (*P* < 0.05). The early flowering phenotype observed in ZmSOC1-CX lines suggests that the expression of *SlMBP18* and the fifth *SOC1*-like gene of tomato may play a role in promoting flowering. For the T_1_ generation, it would have been valuable to investigate additional traits such as seed germination time and seedling growth, as previous studies have shown that repressed expression of DWARF genes can delay seed germination ^42^. Notably, while GmSOC1-CX has been reported to induce early flowering in *Arabidopsis* (Zhong et al., 2012), its overexpression did not significantly promote flowering in soybean. However, *GmSOC1* knock-out mutants exhibited delayed flowering ^57^, highlighting the complex and species-specific regulatory roles of *GmSOC1* in flowering time control.

Interestingly, in this study, the T_0_ generation of VcSOC1K-CX plants from four transgenic lines exhibited earlier flowering compared to the non-transgenic lines, whereas the T_1_ generation of three transgenic lines did not show this early flowering phenotype. Previous studies have reported that overexpression of *VcSOC1K* in blueberry and *VcSOC1K-CX* in tobacco promoted flowering ^23,58^; however, *VcSOC1K-CX* did not significantly induce early flowering in maize ^24^. This variation may be attributed to many factors, including differences in expression levels, plant species, and genotype.

At the transcript levels, GmSOC1-CX did not promote flowering, at least not significantly, is likely due to the enhanced expression of two Histone-lysine N-methyltransferase genes, *CLF* and *EZA1* (Fig. 5) ^35–39^.

### Plant architecture and the expression of full-length *SOC1* and partial *SOC1*

Plant architecture, including shoot and inflorescence structure, is an omnigenic trait that can significantly influence tomato productivity ^59,60^. Tomato *SOC1* genes, such as *SlTM3* and *SlSTM3*, serve as core regulators of inflorescence structure by interacting with other MADS-box genes ^31,32,59,60^. Overexpression of *SlTM3* and *SlSTM3* enhances inflorescence branching ^31^, while their suppression reduces branching ^32,60^. In this study, transgenic lines expressing *GmSOC1*, *ZmSOC1*, and *VcSOC1K* exhibited no noticeable changes in inflorescence structure. For GmSOC1-CX lines, this was further supported at the transcript level by the unchanged expression of tomato *FRUITFULL1* (*FUL1*)(Table 1), a direct activation target of *SlSTM3* ^31^.

Phenotypically, the effect of *SOC1* expression on plant height and branching has been less consistent in the literature compared to its well-established role in flowering time. In soybean, *soc1* mutants have more internodes than the wild type, but the impact of *GmSOC1* overexpression on plant architecture remains unclear ^57^. In *Medicago truncatula*, *MtSOC1* has been shown to influence both flowering and primary stem height in both mutant and overexpression lines ^61^. Additionally, ZmSOC1-CX expression has been associated with reduced plant height in transgenic maize and soybean plants ^22,33^. In tomato, *SlTM3* and *SlSTM3* expression have not displayed any significant impact on plant height and branching ^32^.

In this study, GmSOC1-CX, ZmSOC1-CX, and VcSOC1-CX did not significantly change plant height but enhanced branching for at least the GmSOC1-CX and VcSOC1-CX plants (Fig. 2A-C).

At the transcript level, it is noteworthy that the repressed expression of four DWARF genes could theoretically lead to reducing plant size in tomato due to less BRs production, as suggested by previous studies (Fig. 5) ^41–43,62,63^. The enhanced expression of *CKX7* could result in reduced cytokinin levels, which are expected to shorten plant height, increase branching, and reduce flower number ^48,64–66^. Similarly, a decreased expression of *KAO2* might reduce GA production ^49^, potentially leading to delayed seed germination, stunted plant growth, and delayed flowering. In this study, although GmSOC1-CX did not exhibit phenotypic changes such as plant dwarfing, delayed flowering, or reduced flower number, it did show increased branching (Fig 1C-F, Fig. 2A-C, and Supplementary Fig. S1D). However, BR, cytokinin, and GA levels, as well as seed germination timing, were not investigated in this work.

### Tomato fruit yield and the expression of full-length *SOC1* and partial *SOC1*

Crop yield-defining traits vary across different crops ^27^. For tomatoes, yield-defining traits include both direct factors, such as fruit number, fruit size, and fruit production efficiency per unit area, as well as related traits, including tolerance to abiotic and biotic stresses ^26,60^.

Accordingly, hormone and flowering pathway genes have become the targets for genetic improvement of yield ^29–31,67,68^. In this study, two full-length *SOC1* genes from maize and soybean, along with the *K*-domain of the blueberry *SOC1* gene, were constitutively expressed in tomato. The resulting transgenic lines exhibited an increased fruit count, leading to higher total fruit production per plant, suggesting enhanced yield potential. This rise in fruit number was associated with improved branching in the transgenic plants, likely due to decreased BRs resulting from the repression of *DWARF* genes. Notably, similar effects have not been reported in tomato through the overexpression of *SlTM3* and *SlSTM3*. Interestingly, lower expression levels of *GmSOC1* have been shown to enhance soybean yield ^57^.

In addition to its essential role in flowering, *SOC1* also plays a role in other processes in *Arabidopsis*, such as root development and leaf senescence, both of which impact crop yield ^69,70^. However, the impact of *SOC1* overexpression on root development and leaf senescence in crops remains largely unexplored. Previously, we found that constitutive expression of three *SOC1* genes, either full-length or partial, has the potential to enhance yield in maize, soybean, and blueberry, primarily through the regulation of flowering pathway genes, as indicated by RNA-seq data ^22–24,33^. In this study, unlike our previous findings, the observed increase in fruit production per transgenic plant is attributed to enhanced branching, likely due to the repressed expression of *DWARF* genes (Fig. 5). Additionally, the increased expression of *CLF* and *EZA1* in the RNA-seq data of the *GmSOC1-CX* lines appears to provide evidence explaining the unchanged flowering time (Fig. 5).

### Tomato fruit quality, biotic and abiotic tolerance, and other traits

In this study, the biotic and abiotic tolerance of the transgenic plants were not directly assessed. However, the identification of several DEGs associated with these traits suggests that the transgenes may have influenced plant resilience to biotic and abiotic stresses (Fig. 4, Table 1). Additionally, fruit quality as well as the other traits may have been impacted by the DEGs.

## Materials and methods

### Constructs and plant transformation

The ZmSOC1 and GmSOC1 protein sequences were used as queries to conduct BLAST searches against the tomato proteome in Phytozome v13. The identified tomato SOC1 and SOC1-like proteins were subsequently selected for phylogenetic analysis (Fig. 1A).

A 696-bp *ZmSOC1* (also known as *ZmMADS1*), identical to the sequence derived from HQ858775.1 in GenBank, was cloned using polymerase chain reaction (PCR) from the cDNA of the maize inbred line B104 ^22^. The corresponding protein sequence matches NP_001105152.1 in GenBank (Fig. 1A). The fragment was inserted into a modified pCAMBIA1300 vector between a CaMV 35S promoter and a CaMV poly(A) signal at the *KpnI* and *XbaI* restriction sites (Fig. 1B).

Similarly, a 630-bp *GmSOC1*, identical to the sequence derived from NM_0011249448.2 in GenBank, was cloned from the cDNA of the soybean cultivar Thorne. The fragment was inserted into the same modified pCAMBIA1300 vector between a CaMV 35S promoter and a CaMV poly(A) signal at the *KpnI* and *XbaI* sites (Fig. 1B). The corresponding protein sequence is identical to NP_001236377.1 in GenBank (Fig. 1A).

The *VcSOC1K* gene was previously cloned into the T-DNA region of the binary vector pBI121, positioned between the CaMV 35S promoter and the Nos terminator for constitutive expression ^23^. In this study, the *VcSOC1K* expression cassette was excised from pBI121 using *Hind*III and *EcoR*I digestion and subsequently ligated into the T-DNA region of a *HindIII*- and *EcoR*I-digested pCAMBIA1300 (PC1300) vector. This ensures that all three constructs share the same binary vector backbone (Fig. 1B).

All three constructs were confirmed by sequencing the target genes and subsequently transformed into *Agrobacterium tumefaciens* strain EHA105. For tomato transformation, indeterminate tomato (*Lycopersicon esculentum*) ‘Ailsa Craig’ was used. Seeds were sterilized in a 2.5% (v/v) sodium hypochlorite solution and germinated to produce cotyledons. Non-inoculated cotyledons were cultured on regeneration medium without antibiotics to generate non-transgenic (NT) regenerants, which served as controls. Shoots approximately 2-3 cm in length were excised from the regenerated explants and transferred to 30 mL Murashige and Skoog (MS) medium for rooting ^71^. The rooting medium was either free of antibiotics for NT plants or contained 10 mg/L hygromycin, 250 mg/L timentin, and 250 mg/L cefotaxime.

### Transplanting, phenotyping, and genotyping of the transformants

After rooting, T_0_ plantlets were transplanted into 4-inch plastic pots containing water-soaked Suremix Perlite planting medium (Michigan Grower Products Inc., Galesburg, MI). The plants were covered with plastic bags and placed in a growth room maintained at 25 °C with a 16/8- hour light/dark photoperiod. Over a two-week period, the plastic bags were gradually removed to acclimate the plantlets. Once acclimated, the plants were repotted into one-gallon pots and transferred to a greenhouse.

For each construct, approximately 10-20 T_0_ transformants were cultivated in the greenhouse. Each transformant, derived from a separate explant, was considered an independent transgenic line. At the time of repotting into one-gallon pots, transformed and non-transformed plants of similar size, 3-5 lines per construct and non-transformants, were selected. These comparable plants were used as representatives for phenotyping of T_0_ plants. First-generation (T) seeds were harvested separately from each T_0_ plant.

For phenotyping T_1_ plants, 20-30 seeds from each of three selected T_0_ transgenic lines per construct were germinated in soil. Ten plants from each transgenic line were transferred to one-gallon pots and grown in a greenhouse.

Phenotypic assessments for each plant included measuring the time and height of the first flower appearance, recording the time of the first mature fruit appearance, counting the total number of branches and fruits harvested, and weighing all harvested fruits. Photographs were taken to document phenotypic variations.

For genotyping of the T_0_ and T_1_ plants, genomic DNA was extracted from young leaves using the cetyltrimethylammonium bromide (CTAB) method ^72^. PCR was performed to detect the *hph* transgene and the full-length sequences of *GmSOC1*, *ZmSOC1*, and *VcSOC1K* using the primers listed in Table S1.

### RNA sequencing and quantitative reverse transcription PCR analysis

Three groups of T_1_ plants, derived from three T_0_ transgenic lines containing *GmSOC1*, were selected for RNA sequencing and transcriptome analysis. Each group of T_1_ plants was segregated into transgenic and NT plants. For each group, two newly formed mature leaves from all transgenic plants before their flowering were pooled together to create one biological replicate for transgenic plants, while leaves from NT plants were similarly pooled to form one biological replicate for NT plants. In total, three transgenic samples and three NT samples were collected, immediately frozen in liquid nitrogen, and stored at -80°C for RNA isolation.

Total RNA from each sample was isolated using a modified CTAB method ^73^ and further purified with the RNeasy Mini Kit (Qiagen, Valencia, CA, United States). To eliminate any residual DNA, on-column DNase digestion was performed using the RNase-Free DNase Set (Qiagen, Valencia, CA, United States). RNA quality was assessed with the High Sensitivity RNA ScreenTape system (Agilent Technologies, Santa Clara, CA, United States). All RNA samples used for sequencing and RT-qPCR analysis had RNA integrity number equivalent scores exceeding 5.0.

The RNA samples were sequenced using the Illumina NovaSeq 6000 platform (150 bp paired-end reads) at the Research Technology Support Facility at Michigan State University (East Lansing, Michigan, United States). The quality of the sequencing reads was evaluated using the FastQC program, focusing on per-base quality scores. Each of the six biological samples yielded 19-21 million paired reads (MR), with average quality scores exceeding 35, ensuring suitability for transcriptome analysis.

A transcriptome reference was assembled from approximately 120 million paired reads (MR) obtained from all six NT and transgenic lines using Trinity v2.15.1 ^74^. This reference was used for differential expression analysis. Transcripts identified as differentially expressed (DETs) with a false discovery rate (FDR) below 0.05 were selected for further analysis of various pathway genes. The transcriptome reference was annotated using Trinotate/4.0.2.

Pathway genes for nine phytohormones in *Arabidopsis*, including auxin, cytokinin, abscisic acid (ABA), ethylene, gibberellin (GA), BRs, jasmonic acid, salicylic acid, and strigolactones, were retrieved from the RIKEN Plant Hormone Research Network. Similarly, sugar pathway genes in *Arabidopsis* were identified. These hormone, MADS-box, and sugar pathway genes from *Arabidopsis* were used as queries to perform BLAST searches against the transcriptome reference, and isoforms with e-values less than -20 were identified for transcriptome comparisons. Flowering pathway genes in *Arabidopsis* and cereals ^75^ were used to analyze flowering-related differentially expressed transcripts (DETs) identified in this study. Gene networks of overrepresented gene ontology (GO) terms for the selected DETs were constructed using Cytoscape 3.10.3.

Six selected DEGs were further analyzed through RT-qPCR using using the SYBR Green system (LifeTechnologies, Carlsbad, CA). A tomato *ACTIN* gene served as the reference gene to normalize the RT-qPCR results. All primers used in the analysis are included in Table S1. The same RNA samples used for RNA sequencing, including three biological samples and three technical replicates, were used for the RT-qPCR analysis. Fold changes were calculated using 2^−ΔΔCt^, where ΔΔCt = (CtTARGET – CtNOM)transgenic – (CtTARGET – CtNOM)non-transgenic.

## Author Contributions

GHD: Development of transgenic lines and phenotyping of T0 plants; JJ: RNA-sequencing analysis; GS: Study conception and supervision, data analysis and interpretation, manuscript drafting, and finalization. All authors reviewed the results and approved the final version of the manuscript.

## Data availability

All data analyzed during this study are included in this published article.

## Acknowledgments

We extend our gratitude to Dr. Xue Han for her assistance in caring for the plants in the greenhouse. The work was supported partially by AgBioResearch of Michigan State University. Dr. Gharbia H. Danial’s study in the US were supported by Ministry of the Higer Education in the Kurdistan region in Iraq.

## Supporting Information

**Table S1** PCR and qRT-PCR primers

**Supplementary Fig. S1 Phenotypic comparisons of T_0_ transgenic lines (GmSOC1_CX, *n* = 5; ZmSOC1_CX, *n* = 4; and VcSOC1K_CX, *n* = 3) and their corresponding non-transgenic (NT) lines (NT_GmSOC1, *n* = 4; NT_ZmSOC1, *n* = 3; and NT_VcSOC1K, *n* = 3). (A)** Days to the appearance of the first flower after potting in a one-gallon pot. **(B)** Plant height at the time of first flowering. **(C)** Number of flower clusters counted after all fruits were harvested. **(D)** Number of branches counted after all fruits were harvested. **(E)** Days to the appearance of the first mature fruit after potting in a one-gallon pot. **(F)** Total number of fruits harvested. **(G)** Total weight of harvested fruits. **(H)** Average weight per fruit. The y-axis shows averages, and bars indicate standard deviations. Significance codes: \*\**p* < 0.01, \**p* < 0.05.

