## Supplementary figures and images for "Constitutive expression of full-length or partial of *SOC1* genes for yield enhancement in tomato"

### Figure S1

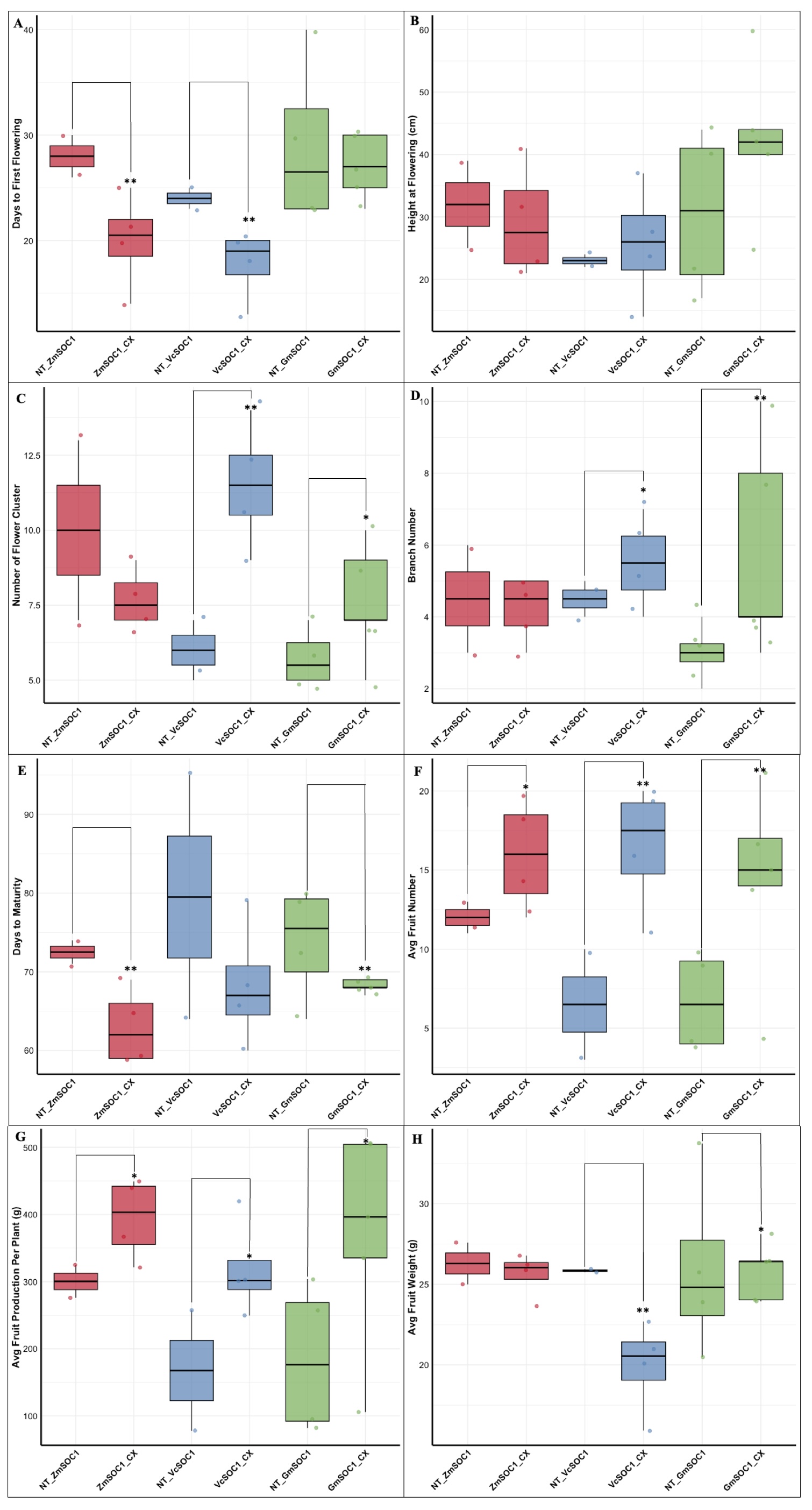
